# The interplay between host biogeography and phylogeny in structuring diversification of the feather louse genus *Penenirmus*

**DOI:** 10.1101/2021.06.21.449287

**Authors:** Kevin P. Johnson, Jason D. Weckstein, Stephany Virrueta Herrera, Jorge Doña

**Affiliations:** Illinois Natural History Survey, Prairie Research Institute, University of Illinois, Champaign, Illinois, USA; Department of Ornithology, Academy of Natural Sciences of Drexel University and Department of Biodiversity, Earth, and Environmental Sciences, Drexel University, Philadelphia, Pennsylvania, USA; Program in Ecology, Evolution, and Conservation, University of Illinois, Urbana, Illinois, USA; Departamento de Biología Animal, Universidad de Granada, Granada, Spain

**Keywords:** parasite, coevolution, host-switching, Phthiraptera, Piciformes, Passeriformes

## Abstract

Parasite diversification is influenced by many of the same factors that affect speciation of free-living organisms, such as biogeographic barriers. However, the ecology and evolution of the host lineage also has a major impact on parasite speciation. Here we explore the interplay between biogeography and host-association on the pattern of diversification in a group of ectoparasitic lice (Insecta: Phthiraptera: *Penenirmus*) that feeds on the feathers of woodpeckers, barbets, and honeyguides (Piciformes) and some songbirds (Passeriformes). We use whole genome sequencing of 41 ingroup and 12 outgroup samples to develop a phylogenomic dataset of DNA sequences from a reference set of 2,395 single copy ortholog genes, for a total of nearly four million aligned base positions. The phylogenetic trees resulting from both concatenated and gene-tree/species-tree coalescent analyses were nearly identical and highly supported. These trees recovered the genus *Penenirmus* as monophyletic and identified several major clades, which tended to be associated with one major host group. However, cophylogenetic analysis revealed that host-switching was a prominent process in the diversification of this group. This host-switching generally occurred within single major biogeographic regions. We did, however, find one case in which it appears that a rare dispersal event by a woodpecker lineage from North America to Africa allowed its associated louse to colonize a woodpecker in Africa, even though the woodpecker lineage from North America never became established there.

## 1. INTRODUCTION

Understanding the diversity of life on Earth requires clear identification of factors governing speciation for an array of different organisms. Although parasites represent a large fraction of all species (de Meeûs and Renaud, 2002), the interplay of forces responsible for parasite speciation remains poorly understood. Parasite speciation is influenced by the same factors that affect free-living organisms, such as biogeography (Thompson, 2005; Sobel et al., 2010). Parasite speciation is also influenced by the ecology and evolution of their hosts. Therefore, integrating these factors is key to understanding parasite diversification (Thompson, 2005; Clayton et al., 2015).

Permanent parasites, which spend their entire lifecycle on the host, are excellent models for studying parasite diversification, because the ecology and evolution of the host lineage can have a major impact on parasite diversification (Clayton et al., 2015). In particular, the close association between host and parasite in these systems can result in cospeciation, i.e. simultaneous divergence of host and parasite lineages (Brooks, 1979). However, even in these tightly interacting systems, host-switching can also be a common process (Johnson et al., 2002; Doña et al., 2017; Boyd et al., In review). Host-switching involves parasite colonization (i.e. successful dispersal and establishment) of a host species on which it did not previously occur (Combes, 2001; Clayton et al., 2015). Normally host-switching requires biogeographic overlap between the involved host species. Thus, biogeographic patterns and processes may also be extremely important in the diversification of parasite lineages (Sweet et al., 2018).

The feather lice (Insecta: Phthiraptera: Ischnocera) of birds, which are permanent parasites, have been an important system in studies of the influence of biogeography on parasite speciation(Clayton et al., 2015). These parasites spend their entire lifecycle among the feathers of their avian hosts; gluing their eggs to feather barbs and molting through three nymphal instars. Most dispersal of feather lice is through vertical transmission, between parents and offspring (Harbison et al., 2008). However, transmission can also occur horizontally through direct contact between individual birds during interactions (Darolova et al., 2001), and in some groups of feather lice through phoresis (hitch-hiking) on hippoboscid flies (Harbison et al., 2009). Other aspects of shared habitat, such as dust baths or hole nests, may also facilitate dispersal by feather lice between birds (Johnson et al., 2002). Previous studies of feather lice (Weckstein, 2004; Sweet and Johnson, 2018; Sweet et al., 2018; Boyd et al., In review) have shown that biogeographic factors facilitating dispersal and switching of lice among host species may be just as important to understanding the diversification of lice, as knowledge of host diversification itself.

In addition to biogeography, host switching may be limited by the physical and morphological features of the host (Clayton et al., 2003). Many genera of feather lice are restricted to a single family or order of hosts (Price et al., 2003), even though they appear to readily switch among different species of hosts in that group (Weckstein, 2004; Boyd et al., In review). Of particular interest in this regard are cases of “major” host switching, i.e. switching of louse lineages between different families and orders of birds (Johnson et al., 2011; Clayton et al., 2015). Some phylogenetic examples of major host switching in feather lice include the switching of wing lice (*Anaticola*) from flamingos (Order: Phoenicopteriformes) to waterfowl (Order: Anseriformes) (Johnson et al., 2006) and switching of body lice between landfowl (Order: Galliformes) and pigeons and doves (Order: Columbiformes) (Johnson et al., 2011).

Here we focus on a single genus of feather louse (*Penenirmus*) that is found on two different avian orders (Piciformes and Passeriformes), with a nearly worldwide distribution (Price et al., 2003). Among the Piciformes, species of *Penenirmus* parasitize several families, including Old World barbets (Megalaimidae and Lybiidae), New World barbets (Capitonidae), honeyguides (Indicatoridae), and woodpeckers (Picidae). Although toucans (Ramphasitidae) are phylogenetically nested within barbets, this avian group, which has been well sampled for ectoparasites (Weckstein, 2004; Hellenthal et al., 2005; Price and Weckstein, 2005; Price et al., 2004), is not is not known to host any species of *Penenirmus*. Most species of Piciformes are hole-nesting (Winkler et al., 1995), which might facilitate major host-switches, given that the same hole can sometimes be used sequentially by different species and that nest hole takeovers often occur (Winkler and Christie, 2002). Among the songbirds (Passeriformes), the host distribution of *Penenirmus* is more patchy. Although species of this genus are recorded from over 10 songbird families (Price et al., 2003), *Penenirmus* is neither as diverse nor as widespread as many other songbird associated generic groups (e.g. *Myrsidea*, *Brueelia-*complex, *Philopterus*-complex). Prior morphological (Carriker, 1963) and molecular phylogenetic (Johnson et al., 2001) studies have indicated that species of *Penenirmus* from songbird hosts may form a monophyletic group, sometimes recognized as the genus *Picophilopterus*. Given that members of *Penenirmus* occur in multiple biogeographic regions, on two orders of birds, and are widespread across multiple families of Piciformes, this genus is a good candidate for studying the interplay between host biogeography and phylogeny on the diversification of parasites.

Here we reconstruct a phylogenomic tree of *Penenirmus*, sampling specimens of this genus from 41 species of hosts across the diversity of major host groups and biogeographic regions in which it occurs. We leverage genome sequencing data to construct a phylogenomic dataset from 2,395 single copy ortholog genes assembled using aTRAM (Johnson et al., 2013; Allen et al., 2015, 2018). We compare the resulting phylogeny of these lice to a phylogeny of their avian hosts and evaluate the relative influence of host biogeography and phylogeny on the diversification of these parasites.

## 2. METHODS

### 2.1. Taxon Sampling

Samples of *Penenirmus* from 41 species of hosts were selected for genomic sequencing (Table 1). We also used information on the higher level phylogenetics of feather lice (de Moya et al., 2019a; de Moya, 2021), to select 12 species of lice from 9 genera as outgroups, with *Vernoniella* selected as the genus on which the phylogenetic analyses were rooted (Table 1). The taxonomy of the genus *Penenirmus* has proven to be extremely complicated. Many species that parasitize woodpeckers were placed in synonymy (Dalgleish, 1972), with extensive morphological overlap even between the two most widespread species, *P. pici* and *P. auritus*. There has never been a comprehensive revision of the genus, and many species would be difficult to identify based on existing morphological descriptions. Previous Sanger DNA sequence data from a limited number of samples (Johnson et al., 2001) also indicated the potential for cryptic species within currently delimited morphospecies. In addition, *Penenirmus* from many of the hosts sampled for our current study represent new host associations with unknown taxonomic status. Thus, for the purposes of this current paper, we applied names to samples based on host associations described by Price et al. (2003), but considered these assignments to be provisional pending further taxonomic revision. We anticipate that the results of the current study will help inform any future morphologically based classifications.

**Table 1.** Samples in study.

| **Genus** | **species** | **Host** | **Country** | **# Reads** | **# Loci** | **NCBI SRA** |
| --- | --- | --- | --- | --- | --- | --- |
| *Penenirmus* | *auritus* | *Melanerpes aurifrons* | USA | 58889562 | 2327 | SRR9693837 |
| *Penenirmus* | *sp.* | *Melanerpes rubricapillus* | Panama | 79073862 | 2336 | SRR9693819 |
| *Penenirmus* | *auritus* | *Dryocopus pileatus* | USA | 108648814 | 2336 | SRR8566315 |
| *Penenirmus* | *auritus* | *Melanerpes erythrocephalus* | USA | 101173836 | 2326 | SRR8582579 |
| *Penenirmus* | *sp.* | *Eubucco richardsoni* | Brazil | 81873656 | 2334 | SRR8566326 |
| *Penenirmus* | *sp.* | *Eubucco versicolor* | Peru | 43855128 | 2282 | SRR9693810 |
| *Penenirmus* | *auritus* | *Melanerpes cruentatus* | Peru | 59797466 | 2328 | SRR9693836 |
| *Penenirmus* | *auritus* | *Picumnus aurifrons* | Brazil | 73763206 | 2305 | SRR9693812 |
| *Penenirmus* | *auritus* | *Chloropicos goertae* | Ghana | 68459054 | 2325 | SRR9693805 |
| *Penenirmus* | *sp.* | *Sphyrapicus nuchalis* | USA | 72712868 | 2333 | SRR9693834 |
| *Penenirmus* | *auritus* | *Sphyrapicus varius* | USA | 58531158 | 2345 | SRR5308137* |
| *Penenirmus* | *auritus* | *Dryobates pubescens* | USA | 80913408 | 2337 | SRR9693833 |
| *Penenirmus* | *arcticus* | *Picoides tridactylus* | Russia | 68752578 | 2320 | SRR9693840 |
| *Penenirmus* | *sp.* | *Dryobates nigriceps* | Peru | 81775060 | 2338 | SRR8582584 |
| *Penenirmus* | *sp.* | *Capito auratus* | Peru | 61273094 | 2324 | SRR9693811 |
| *Penenirmus* | *sp.* | *Capito aurovirens* | Peru | 70748810 | 2332 | SRR9693835 |
| *Penenirmus* | *sp.* | *Capito brunneipectus* | Brazil | 83340596 | 2327 | SRR9693803 |
| *Penenirmus* | *auritus* | *Colaptes punctigula* | Peru | 91276002 | 2335 | SRR8566314 |
| *Penenirmus* | *auritus* | *Piculus flavigula* | Brazil | 65880460 | 2333 | SRR9693813 |
| *Penenirmus* | *auritus* | *Melanerpes candidus* | Bolivia | 80410572 | 2334 | SRR9693807 |
| *Penenirmus* | *auritus* | *Dendrocopos major* | Russia | 97665712 | 2339 | SRR9693839 |
| *Penenirmus* | *pici* | *Picus canus* | Russia | 75353614 | 2332 | SRR9693838 |
| *Penenirmus* | *marginatus* | *Indicator indicator* | Malawi | 98445194 | 2335 | SRR8566317 |
| *Penenirmus* | *sp.* | *Indicator variegatus* | Malawi | 97859892 | 2335 | SRR8566328 |
| *Penenirmus* | *sp.* | *Indicator minor* | Malawi | 89599388 | 2340 | SRR8566327 |
| *Penenirmus* | *sp.* | *Indicator wilcoxi* | Ghana | 75946788 | 2312 | SRR8173272 |
| *Penenirmus* | *sp.* | *Tricholaema leucomelas* | RSA | 87240610 | 2321 | SRR8582580 |
| *Penenirmus* | *zumpti* | *Lybius torquatus* | RSA | 62994248 | 2321 | SRR9693832 |
| *Penenirmus* | *jungens* | *Colaptes auratus* | USA | 103221024 | 2335 | SRR8566316 |
| *Penenirmus* | *sp.* | *Psilopogon chrysopogon* | Malaysia | 111555534 | 2290 | SRR8582581 |
| *Penenirmus* | *sp.* | *Campylorhynchus turdinus* | Brazil | 95194822 | 2316 | SRR8566322 |
| *Penenirmus* | *sp.* | *Psaltiriparus minimus* | Mexico | 102681560 | 2338 | SRR8566330 |
| *Penenirmus* | *sp.* | *Certhia americana* | Mexico | 79706606 | 2334 | SRR8566323 |
| *Penenirmus* | *sp.* | *Bradypterus baboecala* | Malawi | 83248364 | 2330 | SRR8566319 |
| *Penenirmus* | *sp.* | *Anthus lineiventris* | Malawi | 78153488 | 2333 | SRR8566318 |
| *Penenirmus* | *sp.* | *Cisticola rufilata* | Malawi | 94573310 | 2324 | SRR8566324 |
| *Penenirmus* | *guineensis* | *Lybius dubius* | Ghana | 60628574 | 2343 | SRR9693804 |
| *Penenirmus* | *sp.* | *Chloropicus griseocephalus* | Malawi | 86320320 | 2344 | SRR8566325 |
| *Penenirmus* | *sp.* | *Pogoniulus bilineatus* | DRC | 89048532 | 2342 | SRR8582583 |
| *Penenirmus* | *sp.* | *Gymnobucco calvus* | Ghana | 108180478 | 2347 | SRR8145998 |
| *Penenirmus* | *sp.* | *Gymnobucco peli* | Ghana | 56906082 | 2346 | SRR9693841 |
| **<u>Outgroup</u>** |  |  |  |  |  |  |
| *Turnicola* | *sp.* | *Turnix pyrrhorax* | Australia | 50940150 | 2327 | SRR5308379* |
| *Turnicola* | *sp.* | *Turnix varius* | Australia | 103042686 | 2334 | SRR8146019 |
| *Turnicola* | *angustissimus* | *Turnix nigricollis* | Madagascar | 105324266 | 2332 | SRR8146018 |
| *Cuculoecus* | *africanus* | *Chrysococcyx cupreus* | Ghana | 62937116 | 2347 | SRR5308372* |
| *Craspedorrhynchus* | *subhaematopus* | *Accipiter cooperii* | Canada | 95915920 | 2337 | SRR5308371* |
| *Philopterus* | *sp.* | *Cinnyris afra* | Malawi | 83551206 | 2305 | SRR8566329 |
| *Philopterus* | *sp.* | *Tyrannus melancholicus* | Panama | 53357072 | 2334 | SRR5308375* |
| *Aledoecus* | *sp.* | *Halcyon badia* | Ghana | 77411704 | 2339 | SRR5308110* |
| *Alcedoffula* | *alcyonae* | *Ceryle alcyon* | Canada | 41932010 | 2326 | SRR5308368* |
| *Saemundssonina* | *lari* | *Larus novaehollandiae* | Australia | 38719424 | 2318 | SRR5308141* |
| *Ardeiphagus* | *cochlearius* | *Cochlearius cochlearius* | Brazil | 79452682 | 2334 | SRR5308384* |
| *Vernoniella* | *guimaraesi* | *Crotophaga ani* | Panama | 46644520 | 2332 | SRR5308380* |

\* previously published

### 2.2. Genomic Sequencing

Some of the genome sequencing reads we analyzed here have been previously published (see Table 1 for details). For samples newly sequenced for this study, lice were originally stored in 95% ethanol at −80°C. A single louse was selected for extraction and photographed as a voucher. Total genomic DNA was extracted from this specimen by first letting the ethanol evaporate and then grinding the louse with a plastic pestle in a 1.5 ml tube. A Qiagen QIAamp DNA Micro Kit (Qiagen, Valencia, CA, USA) was used for extraction, and initial incubation at 55°C in buffer ATL with proteinase *K* was conducted for 48 hours. Otherwise, manufacturer’s protocols were followed, and purified DNA was eluted off the filter in a final volume of 50ul buffer AE. Total DNA was quantified using a Qubit 2.0 Fluorometer (Invitrogen, Carlsbad, CA, USA) using the high sensitivity kit.

Genomic libraries were prepared using the Hyper library construction kit (Kapa Biosystems). These libraries were sequenced to generate 150bp paired-end reads using Illumina NovaSeq 6000 with S4 reagents. Libraries were tagged with unique dual-end adaptors and multiplexed at 48 libraries per lane, with a goal of achieving approximately 30−60X coverage of the nuclear genome. Adapters were trimmed and files demultiplexed with bcl2fastq v.2.20 to generate fastq files. Raw reads for each library were deposited in NCBI SRA (Table 1).

### 2.3. Gene Assembly and Phylogenomic Analysis

We used *fastp* v0.20.1 (Chen et al., 2018) to perform adaptor and quality trimming (phred quality >= 30). Trimmed libraries were then converted to aTRAM 2.0 (Allen et al., 2018) blast databases using the atram_preprocessor.py command of aTRAM v2.3.4. We used a reference set of 2,395 single-copy ortholog protein-coding genes from the human louse, *Pediculus humanus*. This reference set has been used in prior phylogenomic studies of hemipteroid insects (Johnson et al., 2018) and the insect order Psocodea (which includes bark lice and parasitic lice, de Moya et al., 2021), and within the *Bemisia tabaci* complex of whiteflies (de Moya et al., 2019b). Thus, this gene set has phylogenetic utility across a wide range of taxonomic scales. The aTRAM assemblies (atram.py command) were conducted using *tblastn* with the amino acid sequences of these genes and the ABySS assembler with the following parameters (iterations=3, max-target-seqs=3000). Exon sequences from these protein-coding genes were then stitched together using the Exonerate (Slater and Birney, 2005) pipeline in aTRAM (atram_stitcher.py command).

The DNA sequences from each sample for each gene were then concatenated together using a custom R script (36 genes that contained sequences from less than 4 samples were discarded at this stage). The nucleotide sequences were translated to amino acids using a custom Python script and aligned based on amino acid sequences using MAFFT v7.471 with the following parameters (--auto --preservecase --adjustdirection --amino) (Katoh et al., 2002, 2013). These aligned amino acid sequences were then back-translated to DNA sequences using the same Python script. Aligned gene sequences were trimmed using trimAL v1.4.rev22 (Capella-Gutiérrez et al., 2009) with a 0.4 % gap threshold. These gene alignments were then concatenated into a supermatrix for phylogenomic analyses using the concat command of AMAS v1.0 (Borowiec, 2016).

A phylogenomic analysis of the concatenated data set under maximum likelihood (ML) was conducted using IQ-TREE 2 v2.1.2 (Minh et al., 2020). We used the -p (Chernomor et al., 2016), −m TESTNEWMERGE (Kalyaanamoorthy et al., 2017), and -rclusterf 10 (Lanfear et al., 2016) parameters to search for the optimal number of partitions and optimal model while maximizing computational efficiency. These parameters were then used in a search using the IQ-TREE algorithm (Nguyen et al., 2015). Tree support was estimated using ultrafast bootstrapping with UFBoot2 (Minh et al., 2013; Hoang et al., 2017).

Because incomplete lineage sorting (ILS) can sometimes result in gene trees that are incompatible with the species tree (Degnan and Rosenberg, 2009), we also used individual gene trees in a coalescent analysis. Individual gene trees were computed under maximum likelihood based on the optimal models using IQ-TREE 2 (-m MFP). These gene trees were then used in a coalescent species tree analysis in ASTRAL-III (Zhang et al., 2018). This software was also used to compute local posterior probabilities for each node in the coalescent tree.

### 2.4. Mitochondrial COI Analysis

Prior molecular phylogenetic study of the genus *Penenirmus* included Sanger sequences of the mitochondrial cytochrome c oxidase I (COI) gene. Thus, for the purposes of comparison of sequences from prior studies, in the current phylogenomic study, we used the same genomic sequence libraries used for assembling nuclear gene sequences above to assemble sequences for the mitochondrial COI gene. In addition, because the mitochondrion is haploid, mitochondrial genes generally sort faster than nuclear genes (Moore, 1995) and have a higher substitution rate in insects, including lice (Johnson et al., 2003a), making them an important tool for understanding patterns of population and species divergence. Understanding the nature of terminal taxa (e.g. species) in a phylogenetic tree is also important for interpreting cophylogenetic analyses, which compare phylogenies of two different lineages (here birds and lice) such that there must be some equivalence in the terminal taxa.

Because sequence reads from the mitochondrion occur in extremely high coverage in these Illumina raw read datasets (generally > 1000X), we used Seqtk v 1.3 (https://github.com/lh3/seqtk) to subsample four million total reads (two million read1 and two million read2) from each library to avoid assembling errors or contaminants. We used a previously published partial COI sequence from *Penenirmus zumpti* (Johnson et al., 2001) in an aTRAM 2.0 (Allen et al., 2018) assembly of the subsampled library from the same species in our study. We ran aTRAM (ABySS assembler, three iterations) to extend the sequence to include the full COI gene. We annotated this sequence based on open reading frames and comparison to *Pediculus humanus*. This new full-length COI sequence was then used as the reference target for assembling COI sequences from all samples in our current study. For these assemblies, aTRAM was run for only a single iteration since we were starting with a full-length sequence as the target. Similar to the approach for the nuclear sequences, COI DNA sequences were translated to amino acids, aligned, and then back-translated to DNA sequences. We blasted COI sequences against NCBI to identify any that were identical, or nearly identical, to previously generated Sanger sequences. For one louse sample with an extremely anomalous biogeographic distribution with respect to its phylogenetic position (from *Chloropicos goertae*, see Results), we obtained an additional louse specimen from the original vial and used Sanger sequencing of a portion of COI following methods of Johnson et al. (2001) to confirm whether this additional sample had an identical (or highly similar) COI sequence to what we obtained from our genomic analysis.

A phylogenetic tree based on these COI sequences was estimated under maximum likelihood using model parameters estimated by IQ-TREE 2. Bootstrap proportions were estimated using ultrafast bootstrapping with UFBoot2. In addition to a tree, we also computed the percent pairwise sequence divergences among all the COI sequences (using the R function *dist.dna*, model “raw”, pairwise.deletion=T from APE v5.5, Paradis and Schliep, 2018) and examined their distribution to provide insights into potential cryptic species or the possibility for future species delimitation.

### 2.5. Molecular Dating Analysis

An estimate of the timeframe of diversification in *Penenirmus* is useful both for comparison with the timing of host diversification and necessary for the biogeographic reconstruction methods employed (below). Because there are no currently known fossilized lice within Ischnocera, to provide calibration points for molecular dating, we use a combination of dates for relevant nodes from prior studies (which typically can calibrate deep nodes in the tree) with terminal cospeciation events (which typically can calibrate shallow nodes).

For deeper calibration points, we used the dating results from an analysis of all nucleotide sites from de Moya (2021), because our current study is also based on an analysis of all sites. The most relevant node to our cophylogenetic analysis (below) is the first split within the focal group (i.e. *Penenirmus*), because we evaluate the ancestral host associated with this node (i.e. the ancestral host of *Penenirmus*). To avoid constraining the date on this node, we did not include any calibration points for this ancestral node, nor the nodes directly above or below it. For nodes present in our tree that were also dated by de Moya (2021), we used the 95% confidence intervals (rounded to the nearest 0.5 mya) as calibrations for these nodes (Supplemental Table S1). For the root of the entire tree, we used the maximum value of the 95% confidence interval as the maximum age for this node (32.0 mya). For more terminal calibration points, we identified nodes in the resulting trees that unite terminal sister species of lice found parasitizing terminal congeneric sister species of hosts. There was only one case of this in our study (*Eubucco richardsoni* versus *versicolor*), and we used the 95% confidence interval from a dating analysis of these birds (Supplemental Table S1, Ostrow et al., pers. comm.) and applied it to the inferred codivergence event in the louse tree.

With these calibrations and the concatenated data set and tree, we used IQ-TREE to perform a dating analysis using the least square dating (LSD2) method (To et al., 2015). Given the overall high support of the nodes in our tree (100% all nodes) and lack of unresolved branches, we set a minimum branch length constraint (u= 0.01) to avoid collapsing short but informative branches without introducing bias to the time estimates (see https://github.com/tothuhien/lsd2). We also inferred confidence intervals by resampling branch lengths 1000 times.

### 2.6. Biogeographic Reconstruction

To evaluate biogeographic patterns in the louse tree, we used the R package BioGeoBears v1.1.2 (Matzke, 2013, 2014), which tests among a variety of biogeographic models and performs biogeographic reconstruction. This approach requires an ultrametric tree; therefore we used the dated louse tree from the molecular dating analysis described above. This tree was pruned to contain only the focal group species (i.e. *Penenirmus*), because the outgroups were not sampled at a taxonomic density sufficient for biogeographic reconstruction. For biogeographic zones, major host groups typically have widespread distributions across broad continental scales, so we defined regions for the lice as New World, Africa, and Eurasia (which also includes southeast Asia). Species of *Penenirmus* do not have any meaningful geographic distribution east of Wallace’s line (Price et al., 2003); thus, Australasia was not included as a biogeographic region.

BioGeoBears was run for the following models: DEC, DIVALIKE, and BAYAREALIKE. We also ran the analyses for the same models with the extra parameter “J” (i.e., to account for jump dispersal events; Matzke, 2014). We then selected the model with the lowest AIC score and used this model to estimate the maximum likelihood ancestral range.

### 2.7. Cophylogenetic Analysis

To compare host and parasite trees, we used eMPRess v1.0 (Santichaivekin et al., 2020). One advantage of this software is that it summarizes events across equally parsimonious (MPR) cophylogenetic reconstructions. To facilitate comparisons with prior cophylogenetic studies, we used costs of duplication:1, sorting:1, and host-switching:2. This is the cost scheme used by most published cophyogenetic studies of lice, as well as other groups of ectosymbionts (Sweet et al., 2016; Doña et al., 2017; Matthews et al., 2018; de Moya et al., 2019a) because duplication+sorting is given an equal total weight to host-switching as alternative ways of reconstructing conflicting host and parasite nodes. Cospeciation always has a zero cost in cophylogenetic reconstruction techniques.

Cophylogenetic reconstructions were restricted to the focal group (*Penenirmus*) and their hosts. For the host tree, we compiled trees from several sources. For backbone relationships among Piciformes, we used the higher-level tree from Prum et al. (2015). To this tree, we grafted branches following published topologies for woodpeckers (Shakya et al., 2017) and Old World barbets (Moyle, 2004). For the passerine species, we downloaded phylogenetic information for all the species in the focal group from BirdTree (Jetz et al., 2012, 2014), and then extracted the subtree corresponding to the passerines. In particular, we downloaded 1,000 trees from the Hackett et al. (2008) backbone tree (only sequenced species) and then summarized those trees by computing a single 50% majority-rule consensus tree using SumTree v 4.5.1 in DendroPy v4.5.1 (Sukumaran and Holder, 2010) following Rubolini et al. (2015). The resulting subtree was also consistent with the tree from Barker et al. (2004). For honeyguides (Indicatoridae), there is no published phylogenetic study, but mitochondrial sequences for the host species in our data set were available in GenBank. From these sequences, we derived a UPGMA tree using Geneious Prime 2020 v0.2 (https://www.geneious.com). For New World barbets (*Eubucco* and *Capito*), we used the topology published by Armenta et al. (2005), which is also supported by an unpublished UCE phylogenomic analysis (Ostrow et al., pers. comm.). For the parasite tree, we used the phylogeny derived from the concatenated data set and dating analysis (above). Because eMPRess does not allow a parasite species to be associated with multiple host species, we represented each host association in our analyses, even for cases in which a single louse species occurred on more than one host species. However, we interpret the results of the cophylogenetic analysis in light of this fact, and do not interpret cophylogenetic events reconstructed within a parasite species. Rather these widespread parasites could either be the result of cohesion (“failure to speciate”) events or recent host switching events (Johnson et al., 2003b; Clayton et al., 2015).

For a given cost scheme, most large cophylogenetic analyses return multiple solutions (MPRs) of equivalent costs. Within eMPRess, it is possible to cluster this MPR space using the Pairwise Distance Algorithm (Mawhorter and Libeskind-Hadas, 2019), where the distance between two MPRs is the number of events that are found in one MPR or the other but not both. As suggested in the tutorial (https://sites.google.com/g.hmc.edu/empress/), we summarized the MPR space into three clusters and drew a representative median MPR for each cluster. Because median MPRs for these clusters did not necessarily satisfy the condition of weak time-consistency, we increased the number of clusters until we got a solution that met this condition. We then evaluated whether the 95% confidence interval for the first divergence in the common ancestor of *Penenirmus* was consistent with the confidence interval for the host node with which that ancestral louse was associated. For this, we used the 95% confidence intervals for the hosts provided by TimeTree (Kumar et al., 2017). Thus, we required our selected set of MPRs to be compatible with the timing of the divergence of the ancestral host for *Penenirmus* and the estimated age for the common ancestor of *Penenirmus*. This consistency in ancestral host and parasite divergence times will also mean a higher probability that derived nodes are also time compatible. Using this MPR, we calculated the cophylogenetic extinction rate (*Ec*; Doña and Johnson, 2020) using a shiny app (https://jdona.shinyapps.io/extinction/). We also computed the costs for 100 random trees in eMPRess under the same costs scheme, to evaluate whether the cost of the actual reconstruction was significantly lower than that for random trees (i.e. significantly more codivergence than expected by chance).

## 3. RESULTS

### 3.1. Genomic Sequencing

Illumina sequencing of genomic libraries from single lice produced between 44 – 112 million total 150bp reads (Read1 + Read2) per sample (Table 1). Assuming a genome size of 200-300 Mbp for Ischnocera (Baldwin-Brown et al., 2021) this would result in nuclear coverage between 22X and 84X. The GC content of the newly sequenced *Penenirmus* libraries was low, typically between 34-38%. Quality scores were high, with mean quality score for each library above 30 at all read positions.

### 3.2. Gene Assembly and Phylogenomic Analysis

Assemblies of 2,395 single copy ortholog genes using aTRAM 2 (Allen et al., 2018) resulted in assemblies ranging from 2,282 to 2,347 genes depending on sample (Table 1). Following alignment, we retained 2,359 genes for phylogenomic analysis. After trimming, the concatenated alignment consisted of 3,917,571 aligned base positions.

Analysis in IQ-TREE (Minh et al., 2020) identified 217 optimal partitions with separate optimal ML models estimated for each. Tree searches with these parameters resulted in a fully resolved tree with all branches supported by 100% of the ultrafast bootstrap replicates (Figure 1). ASTRAL-III (Zhang et al., 2018) gene coalescent searches based on individual gene trees produced a fully resolved tree, with all but one branch supported by 1.0 local posterior probability. The coalescent tree is nearly identical to the concatenated tree, differing in only two branch arrangements (Figure 1). In the concatenated tree, *Penenirmus auritus* from *Melanerpes candidus* is sister to the lice from *Colaptes punctigula* plus *Piculus flavigula*; whereas in the coalescent tree, this louse is sister to the remainder of the *auritus-*complex excluding these two taxa, although this is supported at only 0.84 local posterior probability. The only other difference is that the concatenated tree places the louse from the African Gray Woodpecker (*Chloropicos goertae*) inside the two lice from North American sapsuckers (*Sphyrapicus*), whereas the ASTRAL coalescent tree places this African louse as sister to the two lice from sapsuckers.

**Figure 1.**
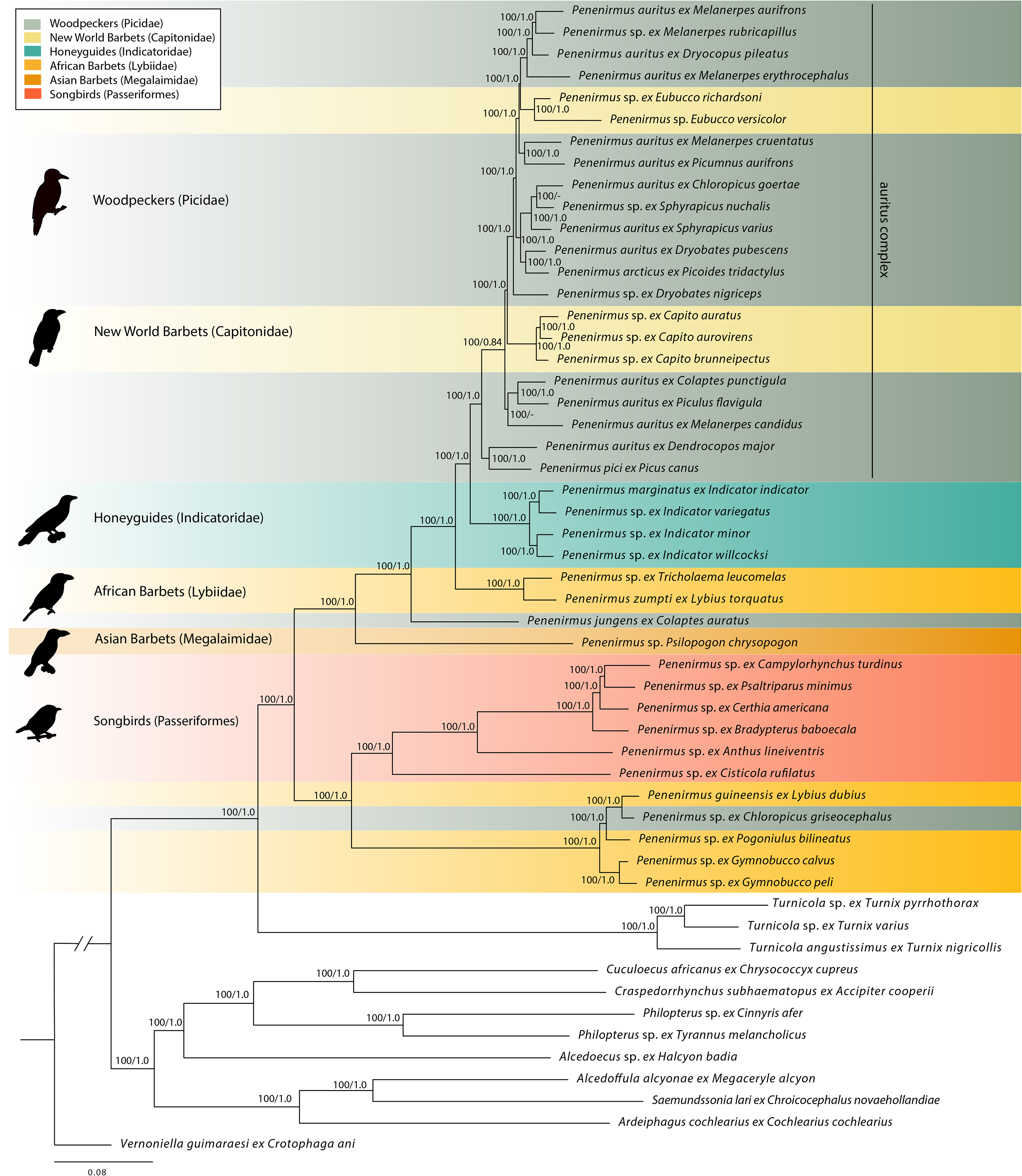
Phylogenomic tree of *Penenirmus* and outgroups resulting from partitioned IQ-TREE ML search of the concatenated data matrix of 2,359 single copy ortholog genes (3,917,571 aligned bp). Branch lengths are proportional to substitutions per site. Numbers on branches are ultrafast bootstraps (from IQ-TREE)/local posterior probabilities (from ASTRAL coalescent analysis). Local posterior probabilities indicated with dash (−) indicate those not recovered by the ASTRAL coalescent analysis (2 branches). Host associations with major groups (Passeriformes and families of Piciformes) are indicated with color shading. Bird silhouettes: Picidae (PhyloPic, Steven Traver), Capitonidae (PhyloPic, Vijay Cavale, John E. McCormack, Michael G. Harvey, Brant C. Faircloth, Nicholas G. Crawford, Travis C. Glenn, Robb T. Brumfield & T. Michael Keesey); Indicatoridae (Wikipedia, Nicolas Huet), Lybiidae (PhyloPic, uncredited), Megalaimidae (Wikipedia, Nicolas Huet), Passeriformes (PhyloPic, uncredited).

Other than these minor differences, these trees provide maximum support (100% ultrafast bootstrap and 1.0 local posterior probability) for many key phylogenetic relationships in *Penenirmus* (Figure 1). The genus *Turnicola* from buttonquails (Turnicidae) is supported as the monophyletic sister taxon of *Penenirmus*, and this is consistent with a study of higher-level feather louse relationships (de Moya et al., 2019) that sampled only a single representative of each genus. In addition, monophyly of the genus *Penenirmus* as currently defined (Price et al., 2003) is recovered. Within *Penenirmus*, two deeply divergent clades are identified. The first contains lice from songbirds (Passeriformes) plus a clade of lice from some African barbets (Lybiidae) and woodpeckers (Picidae). Within this clade, the lice from songbirds form a monophyletic group. The second major clade contains lice from all lineages of Piciformes on which *Penenirmus* occurs, including a second clade of lice from African barbets. Within this second major clade, the one louse sampled from an Asian barbet (Megalaimidae) is sister to the remaining taxa. Other deeply divergent taxa in this second major clade include *Penenirmus jungens* from the Northern Flicker (*Colaptes auratus*), lice from two African barbets (*Lybius* and *Tricholaema*), and a clade of lice from honeyguides (Indicatoridae: *Indicator*). More terminal to these groups is a large clade of lice primarily from woodpeckers (Picidae), but with a few taxa from New World barbets (Capitonidae). Within the clade of woodpecker lice, there is a split between those from Eurasian woodpeckers and those from primarily (though not exclusively) New World taxa. The lice from the two New World barbet genera (*Capito* and *Eubucco*) form separate clades that are not each other’s closest relatives.

### 3.3. Mitochondrial COI Analysis

Assembled sequences of the mitochondrial COI gene were typically highly divergent between species of *Penenirmus,* with most uncorrected pairwise divergences among the ingroup between 20-30% (Supplemental Figure 1). A few comparisons between closely related taxa were on the order of 10% uncorrected pairwise divergence. In addition, there were a few cases of samples differing by less than 5% (all 3.3% or less). These cases of very low divergence (< 3.3%) between louse individuals on different host species included lice on the hosts: 1) *Gymnobucco peli* versus *calva*, 2) *Lybius dubius* versus *Chloropicos griseocephalus*, 3) *Indicator indicator* versus *variegatus*, 4) *Capito auratus* versus *aurovirens*, 5) *Sphyrapicus nuchalis* versus *varius*, and 5) *Dryocopus pileatus*, *Melanerpes aurifrons*, and *M. erythrocephalus*. Thus, most of these cases of highly similar haplotypes on different host species were from hosts of the same genus.

Comparisons of assembled COI sequences to those generated by Sanger sequencing and available in GenBank also revealed cases of highly similar or identical sequences. Six of our assembled COI sequences were identical to those from Sanger sequencing in GenBank. In all cases, these were from the same host species: *Lybius dubius*, *L. torquatus*, *Piculus flavigula*, *Picumnus aurifrons*, *Melanerpes candidus*, and *Chloropicos goertae*. The Sanger sequencing performed for the present study on an individual *Penenirmus* from *Chloropicos goertae* from the same host and vial as our assembled COI sequence from the Illumina reads produced an identical sequence. There were also nine of our assembled COI sequences that had a best Blast hit in GenBank between 0.3% and 2.9% uncorrected sequence divergence. Three of these nine were lice from the same host species. The others were from the same host genus or similar patterns of association that we found with our assembled COI sequences (see above). In one case of note, our assembled COI sequence of the louse from *Gymnobucco peli* was more similar (0.3% divergence) to the GenBank sequence from the louse from *G. calvus*, than our assembled COI of the louse from *G. calvus* was to the Sanger sequence in Genbank of the louse from *G. calvus* (1.3% divergence). All other ingroup assembled COI sequences had best hits in GenBank exceeding 10%.

We did not expect phylogenetic analysis of the assembled COI sequences to produce a highly supported tree, because it is only a single gene with a very high substitution rate in comparison to nuclear loci (Johnson et al., 2003a). However, the tree derived from these sequences (Supplemental Figure 2) in many ways mirrored the trees derived from nuclear loci, particularly for more terminal relationships. In addition, the membership of major clades and the relationships among them were identical to those for nuclear loci (as outlined), except in the case of the COI tree, in which the genus *Turnicola* was embedded within *Penenirmus*. One other relationship of note was that the COI haplotypes of *Penenirmus* lice from the three hosts *Dryocopus pileatus*, *Melanerpes aurifrons*, and *Melanerpes erythrocephalus*, clustered together because of their very low COI sequence divergences, while the louse from *Melanerpes rubricapillus* was sister to this haplotype cluster but differed by approximately 11% from each of them. This was different than the nuclear tree, where the louse from *M. rubricapillus* clustered inside of the clade with these other individual woodpecker lice, perhaps because of ILS in these recently diverged lineages. As expected, the tree generated from COI was much more weakly supported than that from the 2,359 gene nuclear data set.

While many of the samples in our study would traditionally be classified under a single morphospecies, e.g. the widespread *Penenirmus auritus*, these results indicate the presence of highly divergent host-specific haplotype clusters for COI (Supplemental Figure 2). In addition, relationships among terminal taxa derived from COI sequences, particularly within the *Penenirmus auritus*-complex, typically match those from nuclear loci. In one taxon of note, the *Penenirmus* from *Chloropicos goertae*, the relationships from COI match those from the coalescent tree (where *Penenirmus ex Chloropicos goertae* is sister to the lice from the two *Sphyrapicus*), which differs from the concatenated tree. We expect this is because incomplete lineage sorting (ILS) affects the concatenated analysis, but is accounted for by the coalescent analysis. Similarly, the haploid mitochondrial COI gene is also predicted to be less subject to ILS than are nuclear loci (Moore, 1995). Given these considerations, we expect that these clusters of genetically highly similar samples are likely biologically meaningful terminal taxa (i.e. species). Given the distribution of genetic divergences and their host association, we hereafter consider those samples differing by more than 5% uncorrected COI divergence different taxa and those below 5% the same species.

### 3.4. Molecular Dating Analysis

Results of the molecular dating analysis (Figure 2) indicated that *Penenirmus* diverged from its sister taxon (*Turnicola*) at around 23.6 mya (95% CI: 21.2−25.9 mya). The earliest divergence within *Penenirmus* was estimated to be approximately 20.5 mya (18.1 – 22.8 mya). The divergence of the lineage of *Penenirmus* from songbirds (Passeriformes) was estimated to be approximately 17.4 mya (14.9 – 19.8 mya), which is much more recent than the divergence of Passeriformes from other avian orders (62 mya, Prum et al., 2015; or 71 – 86 mya, Kumar et al., 2017). Most of the diversification of the *Penenirmus* from woodpeckers (Picidae) appears to have occurred very recently, with a rapid diversification of lineages in the *auritus*-complex at around 4.0 mya. In general, lineages in the genus *Penenirmus* appear to have diversified well after the major lineages of their hosts, suggesting host-switching may have been an important process in the evolution of host associations for these parasites (see *Cophylogenetics* below).

**Figure 2.**
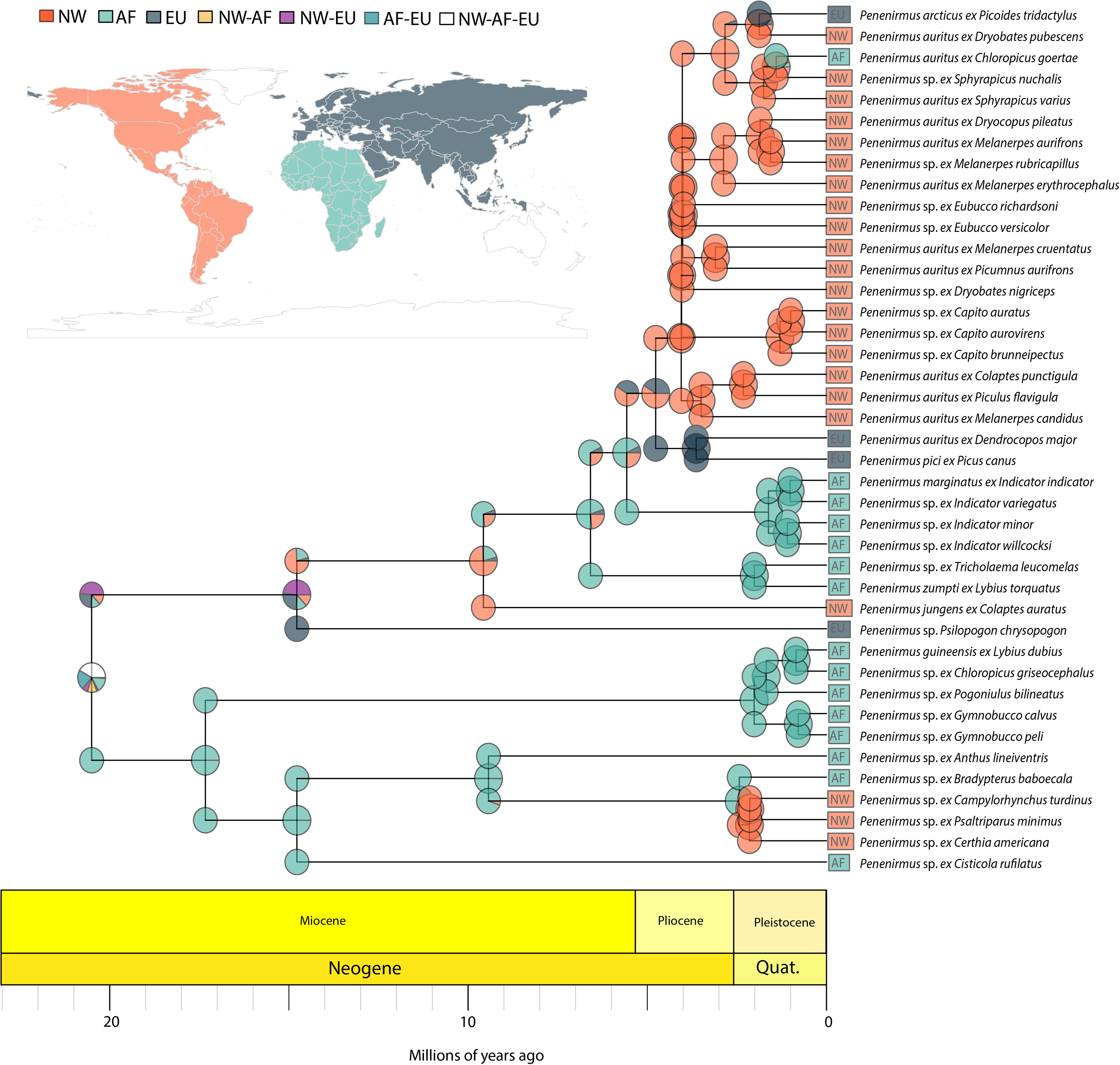
Ultrametric tree of ingroup (*Penenirmus*) resulting from least-square dating analysis including the biogeographic reconstruction from BioGeoBears analysis with major regions color-coded with pie charts proportional to ancestral state likelihoods at ancestral nodes and terminals. Geological timescale indicated at bottom.

### 3.5. Biogeographic Reconstruction

Results of BioGeoBears analysis over the dated ultrametric tree (above) indicated that the DIVALIKE+J (dispersal-vicariance plus long-distance dispersal) was the preferred model with the lowest AIC score (Supplemental Table S2). Maximum likelihood reconstruction within the genus *Penenirmus* under this model (Figure 2) indicated that the geographic distribution of the common ancestor was ambiguous, as was the ancestor of the larger clade parasitizing barbets, honeyguides, and woodpeckers. However, the ancestor of the other major clade (including the clade with songbird hosts) was strongly recovered as having an African distribution, as were most of the lineages in this clade. A dispersal event into the New World from Africa was reconstructed within the lineage of lice parasitizing songbirds, and all of the New World lice in this clade formed a monophyletic group. The ancestral states at some of the deeper nodes within the clade parasitizing barbets, honeyguides, and woodpeckers were less clear. However, at the base of lineages exclusive to a major biogeographic region, such as African barbets, African honeyguides, and Eurasian woodpeckers, these regions were strongly supported at the common ancestors of these lineages. The ancestor of the *P. auritus*-complex was supported as having a New World distribution, with later dispersal by single species into Eurasia (*P. arcticus* on *Picoides tridactylus*) and Africa *(P. auritus* on *Chloropicos goertae*).

### 3.6. Cophylogenetic Analysis

For the cost scheme used in this study, we did not find a median MPR that was at least weakly time-consistent when the number of clusters was set to three or less. When we increased the number of clusters to four, a median weakly time-consistent MPR (i.e., that satisfies the condition that no descendant of a parasite node “p” was mapped to an ancestor of a host node “h”, Santichaivekin et al., 2020) was present in one of the clusters. In this MPR (Figure 3), the ancestral host of *Penenirmus* was inferred by 93% of the reconstructions as the common ancestor of barbets (i.e. Megalaimidae, Lybiidae, and Capitonidae). The 95% confidence intervals for the age of the common ancestor of *Penenirmus* (18.1 – 22.8 mya) and the common ancestor of barbets (22 – 43 mya, Kumar et al., 2017) overlap, making this ancestral host reconstruction compatible with the estimated ages of hosts and parasites.

**Figure 3.**
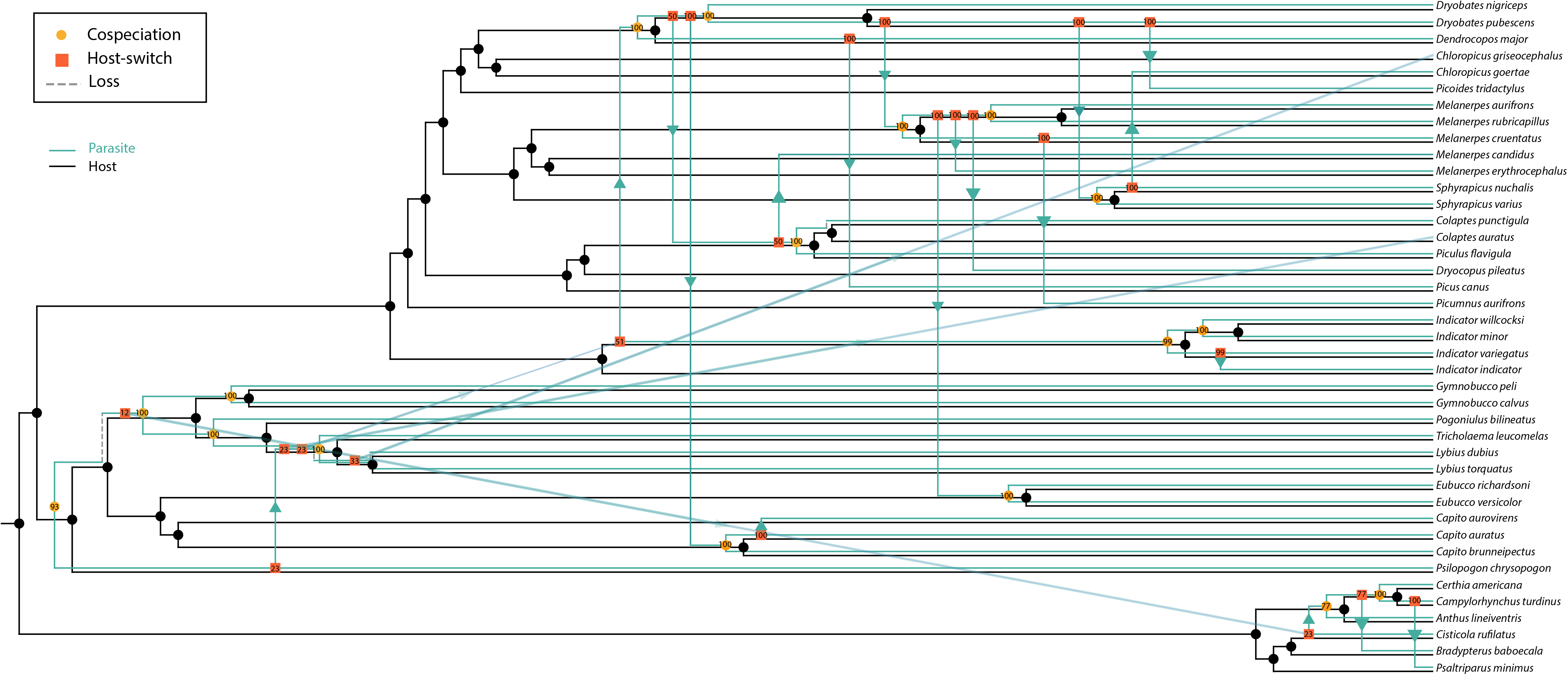
Summary of cophylogenetic reconstruction of optimal MPRs from eMPRess comparison (cost scheme duplication:1, sorting:1, and host-switching:2) of the louse (*Penenirmus*) tree with the avian host tree. Arrows indicate direction of host-switches. Numbers associated with events are the percentage of MPRs with that event.

Cophylogenetic reconstruction in eMPRess that included every sample as a terminal taxon reconstructed 22 codivergence events, 0 duplications, 22 host-switches, and 7 losses (Figure 3). The cost for this reconstruction is much less than that for random trees (*P* < 0.01), indicating more codivergence than expected by chance, even though host-switching is also a prominent process (43% of the events) in the association history of these taxa. The seven reconstructed losses are also relatively high; the cophylogenetic extinction rate (*Ec*) was 0.1 (95% CI: 0.05−0.18), higher than a comparable estimate for avian feather mites (Doña and Johnson, 2020). Considering that some samples represented multiple individuals of the same terminal taxon (using the 5% COI threshold as identified above), five of the host-switches would be interpreted as host-switching with ongoing gene flow (or very recent host-switching) and two would be cohesion events (i.e. failure to speciate by parasites on sister species of hosts).

Songbirds (Passeriformes) were inferred to have acquired their lice from the common ancestor of all African barbets, and this is consistent with Africa reconstructed as the ancestral area for the clade of *Penenirmus* parasitizing songbirds. Honeyguides (Indicatoridae) were also inferred to have acquired their lice via host-switching from a common ancestor parasitizing a more derived lineage of African barbets (*Lybius*). One woodpecker species (*Colaptes auratus*) was also inferred to have acquired its lice via host-switching from this lineage, while a host-switch from the common ancestor of honeyguides to some woodpeckers followed by extensive host-switching within woodpeckers accounts for the distribution of most woodpecker lice. More recent major host-switching events (i.e. between avian families) were also reconstructed, including two host-switches from New World woodpeckers to New World barbets (*Capito* and *Eubucco*), and a host-switch with ongoing gene flow (or very recent host-switch) from an African barbet (*Lybius dubius*) to an African woodpecker (*Chloropicos griseocephalus*). Other than these major host-switching events, the majority of host-switching occurs within a single group (order or family) of birds.

## 4. DISCUSSION

Phylogenomic analyses of sequences targeted from 2,395 single copy nuclear ortholog genes across a broad taxon sampling within the feather louse genus *Penenirmus* produced a very well resolved and highly supported tree. Concatenated ML and coalescent analyses produced nearly identical trees, differing in only two branches, and these trees had all (concatenated) or all but one (coalescent) node supported at 100% (ultrafast bootstrap and local posterior probability, respectively). These trees provide a framework for evaluating biogeographic and host association patterns in this group of avian parasites.

The structure of these trees reflects an interplay between biogeography and host association on the diversification of this genus (*Penenirmus*). Overall, there is considerable biogeographic structure in the parasite tree, with major clades restricted to single biogeographic regions. In part, this is related to the biogeographic distribution of the host groups, but also some of these biogeographically restricted clades occur on multiple major host groups within a region (e.g. barbets and woodpeckers in Africa, woodpeckers and barbets in the New World).

Cophylogenetic reconstructions also support the finding that major host-switches (between families or orders of birds) occur within biogeographic regions and have had important consequences for the diversification of this group. For example, the ancestor of the *Penenirmus* on songbirds is reconstructed as having occurred in Africa, and the cophylogenetic analysis infers that songbirds acquired their lice via a host switch from the common ancestor of all African barbets approximately 17.4 million years ago (mya). Another example of ancient host-switching within a biogeographic region is that from a lineage within African barbets (*Lybius* and *Tricholaema*) to the common ancestor of African honeyguides (*Indicator*) around 6.6 mya. This host-switch was also probably facilitated by the fact that honeyguides are obligate brood parasites (i.e. lay their eggs in the nests of other bird species), and one of the principal avian hosts for these brood parasites are barbets in the genus *Lybius* (Short and Horne, 2002). Thus, honeyguides may have acquired their lice from their foster hosts, as has been shown to sometimes occur for other brood parasites (Lindholm, et al. 1998; Hahn et al., 2000). However, this ancient acquisition appears to have been followed by specialization of these lice on honeyguides and transmission within honeyguide species, since honeyguides are parasitized by a clade of lice restricted to these hosts (Figure 1). This pattern is also the case for the lice of brood parasitic cuckoos (Cuculidae), which sometimes as juveniles possess lice from their foster hosts, but as they age lose these lice and acquire cuckoo-specific lice (Brooke and Nakamura, 1998).

More recent major host-switches also have biogeographic signatures. For example, two lineages of New World barbets (*Capito* and *Eubucco*) were inferred to have independently acquired their lice via host-switching from New World woodpeckers during the last four million years (Figures 2 and 3). Likewise, an African woodpecker (*Chloropicos griseocephalus*) was inferred to have recently acquired its louse via host-switching less than one mya from an African barbet (*Lybius dubius*), perhaps with ongoing gene flow.

The majority of reconstructed host-switching events, however, occur within major host lineages (families or orders). For example, much of the distribution of *Penenirmus* across woodpeckers is inferred to have occurred via host-switching. In part, this is because much of the diversification of woodpeckers occurred before their *Penenirmus* lice (Figure 3). Thus, woodpeckers may have been an open niche for *Penenirmus*, which could have facilitated these host-switches, similar to the case seen in the wing lice *(Columbicola*) of pigeons and doves (Columbiformes; Boyd et al., In review). Hole-nesting behavior within Piciformes may also have facilitated these host-switching events. It is notable that only a single host-switch occurs between Piciformes and Passeriformes, but all others occur within each avian order. None of the songbird (Passeriformes) lice sampled for this study are from hosts that nest in cavities, while all of the Piciformes do nest in cavities (or parasitize the nests of cavity nesting species, in the case of honeyguides). While woodpeckers construct their own nest cavities, many other hole-nesting species rely on naturally occurring cavities or holes constructed by woodpeckers (Winkler and Christie, 2002). Generally, holes for cavity-nesting species are in short supply and thus there can be strong competition between species for these cavities. Interspecific fights and nest cavity take-overs can occur (Winkler and Christie, 2002), which may provide an opportunity for lice to switch hosts, either by physical contact between birds or by lice that were preened off into the nest and remained in the nest at the time of take-over. There are even records of woodpeckers feeding young at the nest of a different species (Winkler and Christie, 2002), which may provide yet another opportunity for louse dispersal between host species.

Other morphological features, such as body size, may also phylogenetically constrain host-switching (Clayton et al., 2003). The species of songbirds sampled here are generally quite small in body size, while most Piciformes are much larger. Louse body size is generally correlated with host body size (Clayton et al., 2003; Johnson et al., 2005), a phenomenon known as Harrison’s Rule (Harrison, 1915). This correlation is likely driven by host preening defenses, with a parasite’s ability to escape from these defenses and remain attached to the host driven by a match between parasite body size and host morphological features, such as the space between feather barbs (Clayton et al., 2003).

While biogeographic distribution of lineages within the genus *Penenirmus* is generally highly conserved (Figure 2), there have been two recent transitions (dispersal events by lice) between major biogeographic regions. The first of these, *P. arcticus*, appears to have been facilitated by host dispersal and speciation. The host of *P. arcticus, Picoides tridactylus*, was inferred to have speciated from its common ancestor with the North American *Picoides dorsalis* by long-distance recent (~2 mya) dispersal across Beringia from North America to Eurasia (Shakya et al., 2017). This date (~2 mya) also matches the split we inferred between the louse *P. arcticus* and its closest relative in the New World.

A more biogeographically enigmatic case is the louse from the African Gray Woodpecker (*Chloropicos goertae*). This louse is very closely related to lice from two North American sapsuckers (*Sphyrapicus varius* and *nuchalis*). While the concatenated analysis actually places the *Penenirmus* from the African Gray Woodpecker as sister to that from the Red-naped Sapsucker *(Sphyrapicus nuchalis*) from western North America, the coalescent and COI analyses place the louse from the African Gray Woodpecker as sister to and divergent from (~10% COI) the lice from the two sapsucker host species. In fact, the COI divergence between the lice from the Red-naped and Yellow-bellied sapsuckers is minimal (<0.5%), suggesting that these lice might be the same species. These two sapsucker species have a broad hybrid zone (Winkler and Christie, 2002), which might provide a mechanism for louse transmission between them, as has been found in mammal lice (Hafner et al., 2019) and feather mites (Doña et al., 2019). The genus *Chloropicos* is not phylogenetically closely related to *Sphyrapicus* (Shakya et al., 2017), and thus neither host phylogeny nor biogeography can explain the very close relationship between the lice from this African woodpecker and those from New World sapsuckers. We also took special effort to assess whether contamination or other lab error could explain these results, and the COI sequences generated via Sanger sequencing of additional specimens from the original field collection vial were identical across three different sequencing attempts of three different louse individuals from this *C. goertae* host sample (recent genome and Sanger sequence this study and the Sanger sequence from Johnson et al. [2001]). While this biogeographic anomaly is difficult to explain, there are records of Yellow-bellied Sapsucker (*S. varius*) from Europe and Atlantic islands (Winkler et al., 1995), which is one of only a few woodpecker species that undergo long-distance migration. Thus, it does seem possible that a wayward ancestral sapsucker may have dispersed to Africa, and while not establishing there, its louse was able to switch to the ancestor of *Chloropicos goertae*. This louse then appears to have become established on this host, perhaps around 1.5 mya, and diverged from its ancestor on sapsuckers. While vagrancy in birds is well documented (e.g. Dunn and Alderfer, 2017), particularly by bird-watchers, this seems to be a case of vagrancy leading to a host-switch and establishment of an avian parasite in a new biogeographic region, even though the original host never became established there. Thus, both species of *Chloropicos* (*griseocephalus* and *goertae*) appear to have acquired *Penenirmus* via host-switching recently (<1.5 mya), which may also be an indication that these woodpeckers were an open niche, perhaps facilitating these host-switches, similar to the case of dove wing lice (Boyd et al., In review).

### 4.1 Taxonomic Implications

Given the lack of recent taxonomic revisions of the genus *Penenirmus*, our results have implications for consideration by future taxonomic revisions. First, the genus *Penenirmus,* as defined by Price et al. (2003), is monophyletic. Some authors (Carriker, 1963) would erect a separate genus (*Picophilopterus*) for the lice on Passeriformes from this group based on morphological differences. While our results certainly support the monophyly of this clade, recognition of this passeriform louse clade at the genus level would render the remainder of *Penenirmus* paraphyletic, because *Picophilopterus* is embedded within *Penenirmus*. Given the morphological distinctiveness and host association of *Picophilopterus*, there might be some merit in recognizing this taxon at the level of subgenus. This would require erecting at least two additional subgenera, perhaps as many as six, for the remainder of the major clades within *Penenirmus*. The number and scope of subgenera would depend on morphological diagnoses as well as maintaining natural groupings that reflect phylogeny.

Our results also have implications for species concepts and delimitation within *Penenirmus*, particularly for *P. auritus* and *P. pici*, the two most widespread species as currently defined (Dalgleish, 1972; Price et al., 2003). The type host for *P. auritus* is *Dendrocopus major*, which is sampled by our study. This sample is sister to *P. pici* from *Picus canus*, which then renders *P. auritus* paraphyletic, and all other lice under this name would need a new species designation under this scenario. We also found that *P. arcticus* from *Picoides tridactylus* was also deeply embedded within *P. auritus*, causing further paraphyly of *P. auritus*. An alternative would be to synonymize *P. pici* and *P. arcticus* into *P. auritus*, which has taxonomic priority, but we do not feel this is warranted given the large genetic divergences, biogeographic patterns, and host associations of the lineages in the phylogenomic tree. In particular, we found that multiple samples from the same host species tended to have nearly identical mitochondrial COI sequences, which were typically highly divergent (>20% uncorrected sequence divergence) from lice from other hosts in the *auritus*-complex. The situation is complicated by the fact that in some cases the lice from different host species are genetically identical, or nearly so, creating a mosaic of patterns of host association and genetic divergence such that heavy reliance on host association alone is not appropriate for full taxonomic revision. Rather, a comprehensive taxonomic revision of the *auritus*-complex is needed with comprehensive taxon sampling and morphological analysis, ideally also incorporating molecular data, to evaluate whether genetic divergences reveal concordant morphological features that might provide a basis for species designation. Given the sweeping synonymy performed by Dalgleish (1972), we suspect that such morphological features exist, but non-overlapping morphological features were difficult to detect in the absence of knowledge of the full scope of host associations of genetically diverged lineages. Thus, further work, both from a molecular and morphological perspective, is needed to understand the species limits and diversification of this prominent and widespread louse genus.

## Acknowledgments

We thank R.J. Adams, A. Aleixo, T. Chesser, R. Faucett, T. Galloway, S. Goodman, A. Gouvea, D. Lane, and C. Witt for assistance in obtaining specimens for this study. We thank A. Hernandez and C. Wright at the University of Illinois Roy J. Carver Biotechnology Center for assistance with Illumina sequencing. Funding was provided by U.S. NSF DEB-1239788, DEB-1925487, DEB-1926919 to K.P.J., NSF DEB-1855812 to J.D.W. and K.P.J., and European Commission grant H2020-MSCA-IF-2019 (INTROSYM: 886532) to J.D.

