## Supplemental figure 1 for "The interplay between host biogeography and phylogeny in structuring diversification of the feather louse genus *Penenirmus*"

***Supplemental Figure 1.*** Plot of pairwise uncorrected percent sequence divergence (y-axis) for each sample (x-axis). Histogram shows relative frequency of sequence divergence categories. Names labeling points are the taxa involved in the comparison with samples labeled on the x-axis.

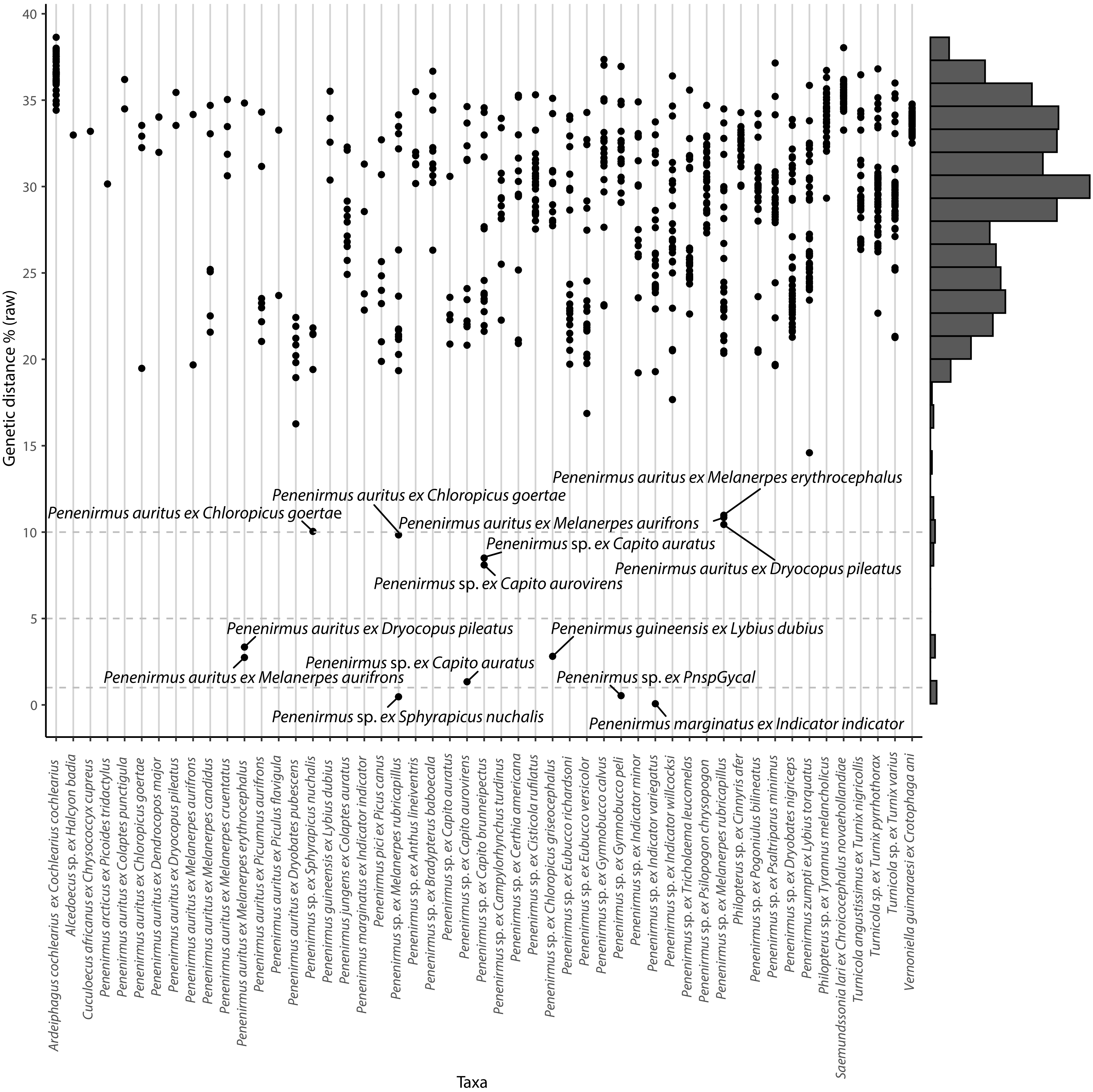
