## Supplemental figure 2 for "The interplay between host biogeography and phylogeny in structuring diversification of the feather louse genus *Penenirmus*"

***Supplemental Figure 2.*** Tree from maximum likelihood analysis of mitochondrial COI sequences. Branch lengths are proportional to subsitutions per site. Brackets denote uncorrected percent sequence divergence between indicated samples. Numbers on branches are ultra-fast bootstrap support values.

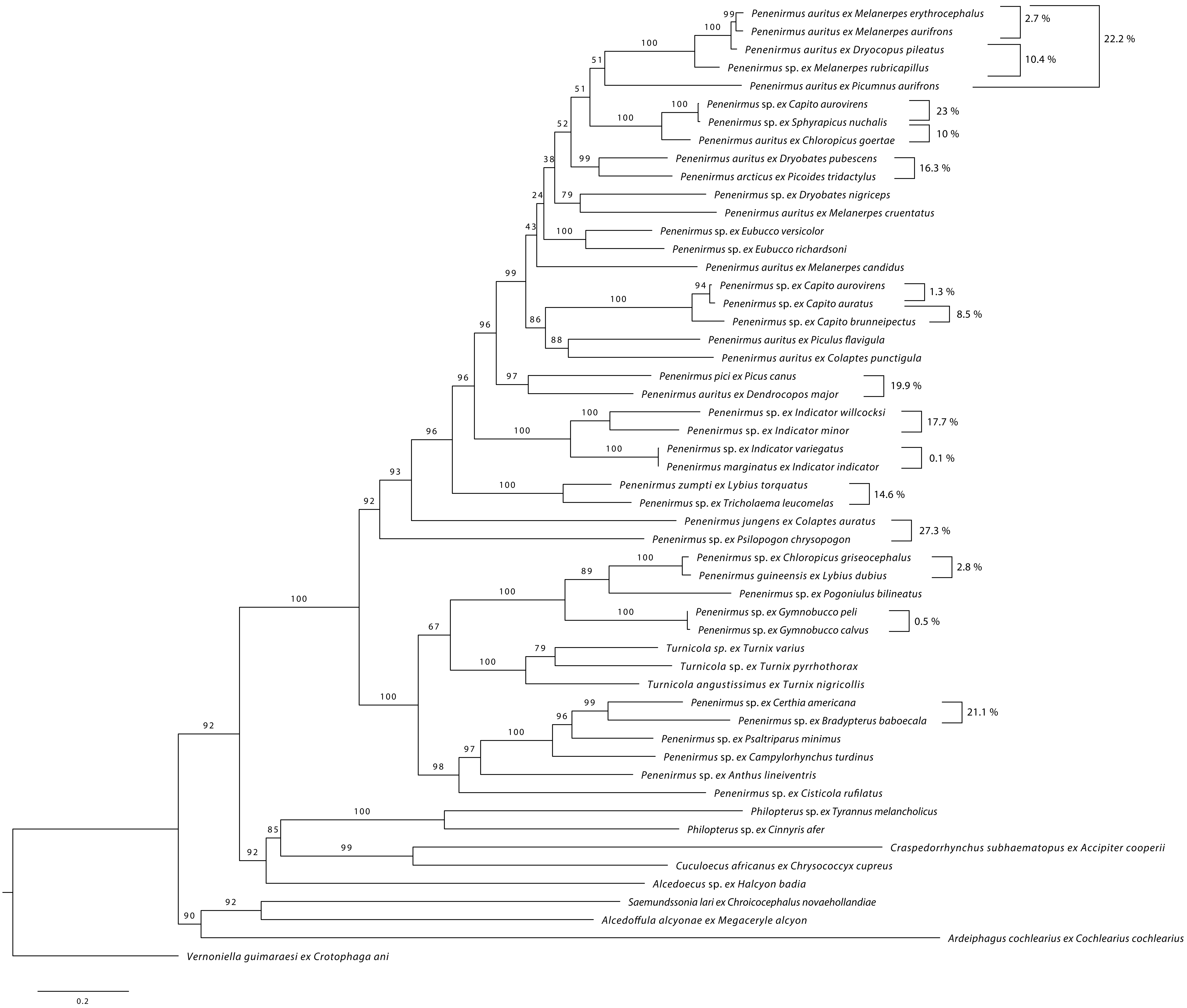
