## Supplementary material for "The interplay between host biogeography and phylogeny in structuring diversification of the feather louse genus *Penenirmus*": Table S1

**Supplemental Table S1. Calibration points used in molecular dating analysis**

| **Calibration Node*** | **Date range** |
| --- | --- |
| *Ardeiphagus cochlearis* | 16.0 – 22.5 mya |
| *Saemundssonia lari* | 12.5 – 17.0 mya |
| *Alcedoecus* sp*.* | 20.0 – 27.0 mya |
| *Philopterus* | 16.5 – 22.5 mya |
| *Craspedorrhynchus subhaematopus* | 9.5 – 16.0 mya |
| *Penenirmus* sp*. ex Eubucco versicolor* | 4.0 – 10.5 mya |

*Indicates date range for node uniting taxon indicated with its sister taxon (from Figure 1)
