## Supplementary material for "The interplay between host biogeography and phylogeny in structuring diversification of the feather louse genus *Penenirmus*": Table S2

**Supplemental Table S2. Model selection for biogeographic reconstruction**

| **Model** | **lnLikelihood** | **Free parameters** | **AIC** |
| --- | --- | --- | --- |
| DEC | -34.00 | 2 | 72.0 |
| DEC + J | -25.64 | 3 | 57.3 |
| DIVALIKE | -32.61 | 2 | 69.2 |
| DIVALIKE + J | -25.29 | 3 | 56.6 |
| BAYAREALIKE | -53.11 | 2 | 110.2 |
| BAYAREALIKE + J | -26.93 | 3 | 59.9 |
